# Functional conservation and divergence of Arabidopsis VENOSA4 and human SAMHD1 in DNA repair

**DOI:** 10.1101/2024.05.27.596045

**Authors:** Raquel Sarmiento-Mañús, Sara Fontcuberta-Cervera, Kensuke Kawade, Akira Oikawa, Hirokazu Tsukaya, Víctor Quesada, José Luis Micol, María Rosa Ponce

**Affiliations:** Instituto de Bioingeniería, Universidad Miguel Hernández, Campus de Elche, 03202 Elche, Spain; Graduate School of Science and Engineering, Saitama University, Saitama City, 338-8570 Saitama, Japan; Center for Sustainable Resource Science, RIKEN, Yokohama, 230-0045 Kanagawa, Japan; Exploratory Research Center on Life and Living Systems, Okazaki, 444-8787 Aichi, Japan; Graduate School of Agriculture, Kyoto University, 606-8502 Kyoto, Japan; Department of Biological Sciences, Graduate School of Science, University of Tokyo, Bunkyo-ku, 113-0033 Tokyo, Japan

**Keywords:** DNA repair, Arabidopsis, *VENOSA4* gene, SAMHD1 ortholog

## Abstract

The human deoxyribonucleoside triphosphatase (dNTPase) Sterile alpha motif and histidine-aspartate domain containing protein 1 (SAMHD1) has a dNTPase-independent role in repairing DNA double-strand breaks (DSBs) by homologous recombination (HR). Here, we show that VENOSA4 (VEN4), the probable *Arabidopsis thaliana* ortholog of SAMHD1, also functions in DSB repair by HR. The *ven4* loss-of-function mutants showed increased DNA ploidy and deregulated DNA repair genes, suggesting DNA damage accumulation. Hydroxyurea, which blocks DNA replication and generates DSBs, induced *VEN4* expression. The *ven4* mutants were hypersensitive to hydroxyurea, with decreased DSB repair by HR. Metabolomic analysis of the strong *ven4-0* mutant revealed depletion of metabolites associated with DNA damage responses. In contrast to SAMHD1, VEN4 showed no evident involvement in preventing R-loop accumulation. Our study thus reveals functional conservation in DNA repair by VEN4 and SAMHD1.

**One sentence summary:** Human SAMHD1 is involved in dNTP metabolism and DNA repair; the latter function is conserved in VEN4, its likely Arabidopsis ortholog.

## INTRODUCTION

In plants and animals, deoxyribonucleoside triphosphate (dNTP) metabolism involves *de novo* biosynthesis, recycling, and degradation. A balanced dNTP pool is essential for maintaining accurate DNA replication and genome stability^1^; its imbalance during DNA biosynthesis can lead to incorrect dNTP incorporations into the nascent strand, which in turn may reduce replication fidelity if mismatched dNTPs are not eliminated, stall DNA polymerases, and induce replication stress.^2–5^ Replication stress affects genome integrity and can be caused not only by dNTP misincorporations but also by the presence of unusual secondary structures in the DNA template strand, or R-loops formed at certain regions by collisions between the transcription and replication machineries (reviewed by Zeman and Cimprich^6^).

Human (*Homo sapiens*) Sterile alpha motif (SAM) and histidine-aspartate (HD) domain-containing protein 1 (SAMHD1) is a nuclear protein with dNTPase activity: it degrades dNTPs into their constituent deoxyribonucleoside and inorganic triphosphate.^7, 8^ SAMHD1 acts in regulating dNTP metabolism and suppressing antiviral immune responses (reviewed by Chen et al.^9^). Moreover, loss of SAMHD1 function has been related to Aicardi-Goutières syndrome, a rare congenital autoimmune disorder that leads to neurodegeneration and typically manifests in early childhood, with symptoms resembling those of a chronic viral infection.^7, 8, 10–12^

Other studies have described a dNTPase-independent role for SAMHD1 in DNA double-strand break (DSB) repair by homologous recombination (HR). SAMHD1 colocalizes with tumor protein p53 binding protein 1 (TP53BP1) at DSB foci produced by treatment with the DSB-inducer camptothecin. At these DSB foci, SAMHD1 recruits C-terminal-binding protein 1-interacting protein (CtIP), an endonuclease that promotes DNA end resection. Cells lacking SAMHD1 function show hypersensitivity to camptothecin, reduced ability to repair DSBs by HR, and impaired recruitment of CtIP to DNA damage foci.^13, 14^ Additionally, in human cell lines, SAMHD1 facilitates the resection of gapped or reversed DNA replication forks by interacting with and stimulating the 3′-5′ exonuclease activity of the DSB repair protein MRE11, which degrades the nascent DNA, thus stabilizing the stalled forks and preventing their collapse into DSBs. Loss of SAMHD1 function also results in the cytoplasmic accumulation of single-stranded DNA fragments from stalled forks, which could induce type-I interferon inflammatory signaling responses.^15^

A recent study demonstrated that SAMHD1 participates in preventing the accumulation of the co-transcriptional R-loops generated in chromatin regions where the transcription and replication machineries collide.^16^ R-loops are three-stranded DNA– RNA structures that form when a nascent RNA anneals with its template DNA strand, displacing the complementary, non-template strand. R-loops are considered a major source of replication stress, DNA breaks, and genome instability, which are a hallmark of cancer; they also impede DNA replication, transcription, and repair (reviewed by Kumar and Remus^17^). Mutations in *SAMHD1* have been identified in different types of cancers and are associated with a poor patient prognosis (reviewed by Li et al.^18^, Mauney and Hollis^19^, Coggins et al.^20^, Chen et al.^21^, and Schott et al.^22^).

We previously performed a large-scale screening for ethyl methanesulfonate (EMS)-induced point mutations causing abnormal leaf shape, size, or pigmentation in Arabidopsis (*Arabidopsis thaliana*). We isolated hundreds of viable mutants, which were classified into 19 phenotypic classes; mutants in the Venosa (Ven) class show reticulation of their rosette leaves.^23–26^ We used map-based cloning to identify the *ven4* mutant, which we refer to as *ven4-0* throughout this paper and in Sarmiento-Mañús et al.^27^ Through iterative linkage analysis of the *ven4-0* mutation using molecular markers, we determined that *VENOSA4* (*VEN4*) is At5g40270, the most likely Arabidopsis ortholog of human SAMHD1.^27, 28^ VEN4 and its rice (*Oryza sativa*) ortholog, STRIPE3 (ST3), are related to chloroplast and leaf development, stress and immune responses, and dNTP metabolism.^27–31^ However, to date, no studies have associated VEN4 or ST3 with DNA damage repair. Here, we analyzed three *ven4* allelic mutants to shed light on whether VEN4 functions in DNA DSB repair by HR. Our results, particularly the study of the original *ven4-0* point mutation, reveal cross-kingdom functional conservation between Arabidopsis VEN4 and human SAMHD1 in DNA damage repair, in addition to their previously known function in dNTP metabolism.

## RESULTS

### Arabidopsis VEN4 and human SAMHD1 share functionally crucial residues

The full-length human SAMHD1 and Arabidopsis VEN4 proteins, comprising 626 and 473 amino acids, respectively, show 37% sequence identity. The structure of human SAMHD1 protein has been extensively studied, and several domains and functionally crucial residues have been identified (reviewed by Morris and Taylor^32^). The SAM domain^33^, a putative protein–protein and protein–nucleic acid interaction domain (reviewed by Qiao and Bowie^34^), is located in the N-terminal region of SAMHD1 (spanning residues 45 to 110^35^) and appears to be necessary for protein stabilization after viral infection^36^; however, VEN4 lacks the SAM domain (Figure 1 and Figure S1).

**Figure 1.**
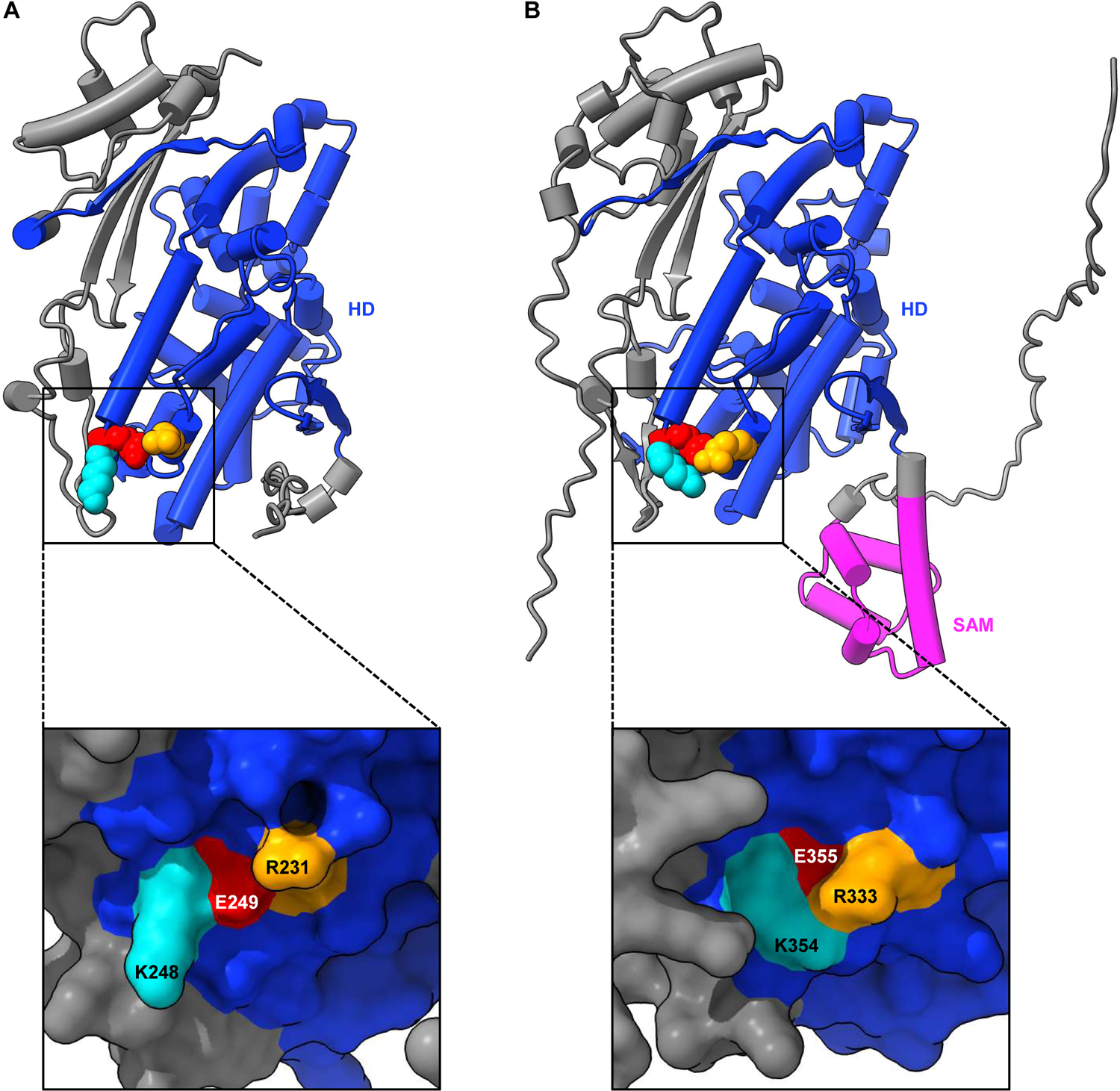
Arabidopsis VEN4 and human SAMHD1 share 3D structural similarity, with conservation of functionally key residues. (A and B) Cartoon and surface representations of the predicted Arabidopsis VEN4 (A) and human SAMHD1 (B) 3D structures. Helices in the cartoon representations are shown as cylinders/stubs. The HD and SAM domains are colored in blue and pink, respectively. The residue altered by the *ven4-0* mutation (E249) and the K248 and R231 residues (which are known to be functionally relevant in SAMHD1) are shown as spheres and highlighted in red, cyan, and orange, respectively (A); the conserved residues in human SAMHD1 are colored similarly (B). Close-up views of the VEN4 and SAMHD1 surfaces have been shaded to highlight protein cavities, which demonstrates that the E249 residue of VEN4 is more accessible to solvent than the equivalent E355 of SAMHD1.

The HD domain^37^ (spanning residues 115 to 465 in human SAMHD1^35^) is highly conserved and required for dNTPase and viral restriction activities.^7, 8, 38–41^ In the HD domain of human SAMHD1, the R451 and L453 residues constitute an RXL motif (where X represents any amino acid). This RXL motif is required for SAMHD1 tetramerization, as well as its dNTPase and retroviral restriction functions, achieved through its interaction with cell cycle regulators such as cyclin A2, cyclin-dependent kinase 1 (CDK1), or CDK2 (Figure S1).^42^ We identified an RXL motif in an equivalent position in VEN4 (R335 and L337) as well as in its plant orthologs, except for tomato (*Solanum lycopersicum*), which has an I residue instead of an L (Figure S2). However, this is a conservative substitution, since I and L are similar aliphatic amino acids. Other key residues of the human SAMHD1 HD domain are conserved in its plant orthologs, including K354 (K248 in VEN4), which is required for DNA DSB repair through HR. Moreover, the deacetylation of K354 by Sirtuin 1 (SIRT1) allows SAMHD1 to function in DSB repair^35^.

The negatively charged E249 residue of VEN4, which is mutated to a nonpolar L amino acid in the *ven4-0* mutant, corresponds to E355 in human SAMHD1 (Figure 1 and Figure S1).^27^ This residue is also E in the tomato ortholog of SAMHD1 and VEN4, but D in the orthologs from the other plant species that we used in our multiple sequence alignment (Figure S2). However, E and D are similar, negatively charged amino acids. Interestingly, the E249 residue of VEN4 is adjacent to K248, the equivalent residue of K354 in human SAMHD1 (Figure 1 and Figure S1)^27^, which is involved in DNA DSB repair by HR^35^, as previously mentioned.

Outside of the HD domain, another conserved residue, T592 (T464 in VEN4), is phosphorylated by the cyclin A2/CDK complex and participates in negative regulation of HIV-1 restriction and MRE11-dependent resection of stalled DNA replication forks.^15, 43, 44^ K484 (K374 in VEN4) is crucial for interaction with CtIP and its recruitment to DSBs to promote DNA end resection by HR.^14, 45^

As expected from their almost identical primary structures (Figure S1), the predicted secondary structures of Arabidopsis VEN4 and human SAMHD1 displayed high three-dimensional conservation, except for their N- and C-terminal regions (Figure 1). In fact, both proteins showed a root mean square deviation (RMSD) of 0.897 Å for 371 pruned (best-matching) atom pairs, while the RMSD across all 457 atom pairs was 7.961 Å.

### *ven4* mutants are hypersensitive to hydroxyurea

Ribonucleotide reductase (RNR) catalyzes the limiting step of the *de novo* dNTP biosynthesis pathway in eukaryotes.^46^ In Arabidopsis, the RNR R1 subunit is encoded by the *RNR1* gene^47, 48^, and the RNR R2 subunit by three paralogous genes: *RNR2A*, *RNR2B*, and *TSO MEANING ‘UGLY’ IN CHINESE 2* (*TSO2*), with *TSO2* being the primary contributor to RNR activity. The rice thermosensitive conditional mutants *virescent3* (*v3*) and *stripe1* (*st1*) are mutated at the *V3* and *ST1* genes, which encode the R1 and R2 subunits of RNR, respectively, and display chlorotic leaves with green stripes.^49^

Rice *v3* and Arabidopsis *rnr2a* mutants are hypersensitive to hydroxyurea, a genotoxic agent that inhibits DNA replication by blocking RNR activity, hence reducing the available dNTP levels. Hydroxyurea also produces stalled DNA replication forks with single-strand breaks (SSBs) and DSBs, which ultimately activate ATAXIA TELANGIECTASIA-MUTATED AND RAD3-RELATED (ATR) for cell-cycle arrest and DNA repair.^49–53^ Indeed, Arabidopsis *atr* mutants also exhibit hypersensitivity to hydroxyurea and other DNA-replication-blocking agents.^51^ Expression of the Arabidopsis *RNR1*, *RNR2A*, and *RNR2B* genes is induced by hydroxyurea and *TSO2* expression is induced by the DSB inducer bleomycin, while the *tso2-1 rnr2a-1* double mutant accumulates DSBs even in the absence of genotoxins.^52, 54^

We grew the *ven4-0*, *ven4-2*, and *ven4-3* mutants (see Methods) on medium supplemented with 1 mM hydroxyurea; all plants were hypersensitive to this DNA-replication-blocking agent, showing severely altered morphology: decreased rosette size and increased leaf reticulation (Figure 2A–J) as well as inhibited primary root growth (Figure 2K and Figure S3) compared with their respective wild types and with their growth in the absence of hydroxyurea. We also observed increased beta-glucuronidase (GUS) enzymatic activity in Col-0 *VEN4pro:GUS* transgenic plants when they were grown on medium containing 1 mM hydroxyurea (Figure 2L–O), indicating that *VEN4* expression is responsive to this compound, similar to *RNR1*, *RNR2A*, and *RNR2B*.^31, 50, 52, 53, 55^ Taken together, these results further support a role for VEN4 in dNTP metabolism, previously proposed by Xu et al.^29^, Lu et al.^31^, and Sarmiento-Mañús et al.^27^, and suggest an additional direct or indirect function in the DNA damage response (DDR).

**Figure 2.**
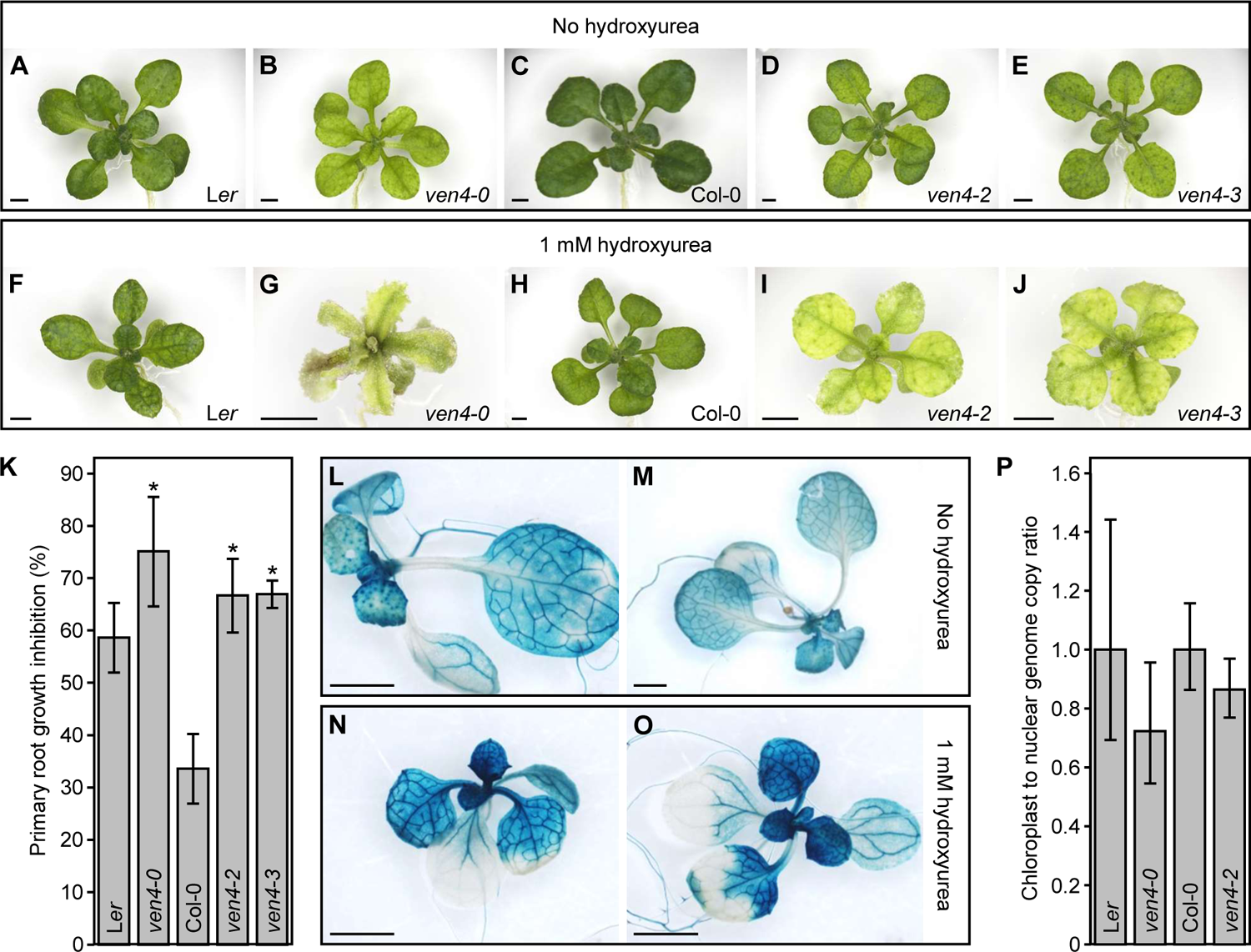
*ven4* mutants are hypersensitive to hydroxyurea but do not show altered chloroplast genome replication. (A–J) Rosettes of the *ven4-0*, *ven4-2*, and *ven4-3* mutants and their L*er* and Col-0 wild types grown on medium not supplemented (A–E) or supplemented with 1 mM hydroxyurea (F–J). Photographs were taken at 19 days after stratification (das). Scale bars: 2 mm. (K) Percentage of root growth inhibition in the *ven4-0*, *ven4-2* and *ven4-3* mutants, and their L*er* and Col-0 wild types, grown for 19 das on vertically oriented Petri dishes containing medium supplemented with 1 mM hydroxyurea. Error bars indicate standard deviation. Primary root length was measured from at least 10 plants. Asterisks indicate values significantly different from the corresponding wild type in a Student’s *t*-test (\**P* < 0.001). (L–O) GUS staining of Col-0 *VEN4pro:GUS* transgenic plants grown on medium not supplemented (L and M) or supplemented with 1 mM hydroxyurea (N and O). Photographs were taken at 11 das. Scale bars: 2 mm. (P) Chloroplast-to-nuclear genome copy ratio in L*er*, *ven4-0*, Col-0, and *ven4-2*, calculated according to Yoo *et al.*^49^ The ratio obtained for the L*er* and Col-0 wild types was set to 1. Error bars indicate the interval delimited by 2^-ΔΔC^T ^± SD^, where SD is the standard deviation of the ΔΔCT values. Three biological replicates were examined per genotype, with three technical replicates in each experiment. The ΔCT values were not significantly different from those of the corresponding wild type in a Student’s *t*-test.

At restrictive growth conditions, the *v3* and *st1* rice mutants display similar chloroplast aberrations to those found in Arabidopsis *ven4* mutants^27, 28^, exhibiting low chlorophyll content and a reduced chloroplast-to-nuclear genome copy ratio.^49^ An explanation has been proposed for these phenotypes: when the dNTP pool is limited, replication of the nuclear genome is likely to be prioritized over that of the chloroplast genome for plant survival.^47, 49^ However, when we studied the chloroplast-to-nuclear genome copy ratio by qPCR amplification of unique sequences from the chloroplast and nuclear genomes in DNA extracted from *ven4-0* and *ven4-2* plants collected 15 days after stratification (das), we did not find a statistically significant reduction compared with their L*er* and Col-0 wild types (Figure 2P).

### DNA DSB repair by HR is impaired in *ven4* mutants

We next aimed to determine whether Arabidopsis VEN4 participates in DNA DSB repair by HR, like its human SAMHD1 ortholog. To this end, we used the *IU.GUS 8* transgene, which was designed for examining somatic HR by the synthesis-dependent strand annealing (SDSA) mechanism. The *IU.GUS 8* transgene contains two overlapping sequences of a non-functional *GUS* gene: a sequence with an internal region of the *GUS* gene inverted (*IU*), and the *GUS* gene lacking this internal region (*GU-US*). In this assay, a GUS-stained blue spot appears as a result of a DSB in the *GU-US* sequence if it is repaired by HR, using the *IU* sequence in the sister chromatid of plants homozygous for the *IU.GUS 8* transgene.^56, 57^

The GUS reporter assay was performed using Hyg^R^ *ven4-0* and *ven4-2* F3 plants (the *IU.GUS 8* construct confers resistance to the hygromycin B antibiotic) derived from crosses of *ven4-0*, and *ven4-2* to the *IU.GUS 8* transgenic line (all in the Col-0 background), which were grown in the presence or absence of 1 mM hydroxyurea as an indirect inducer of DNA DSBs. In the presence of hydroxyurea, we observed on average 1.37 and 0.16 blue sectors per plant in *ven4-0* and *ven4-2* homozygous plants, respectively, and 3.77 and 1.16 in their phenotypically wild-type (*VEN4/VEN4* or *VEN4/ven4*) siblings (Figure 3A–C). We observed 63.76% fewer blue sectors in *ven4-0*, compared with the L*er*/Col-0 hybrid, and 86.2% fewer blue sectors in *ven4-2* compared with their phenotypically wild-type siblings after inducing DNA damage with hydroxyurea. In the absence of hydroxyurea, these mutants also showed fewer blue sectors (34.7% fewer in *ven4-0* and 75.0% fewer in *ven4-2*) (Figure 3A–C). These observations indicate that repair of both spontaneous and hydroxyurea-induced DSBs is impaired in the *ven4* mutants. These results support the hypothesis that VEN4 plays a role in DNA DSB repair by HR.

**Figure 3.**
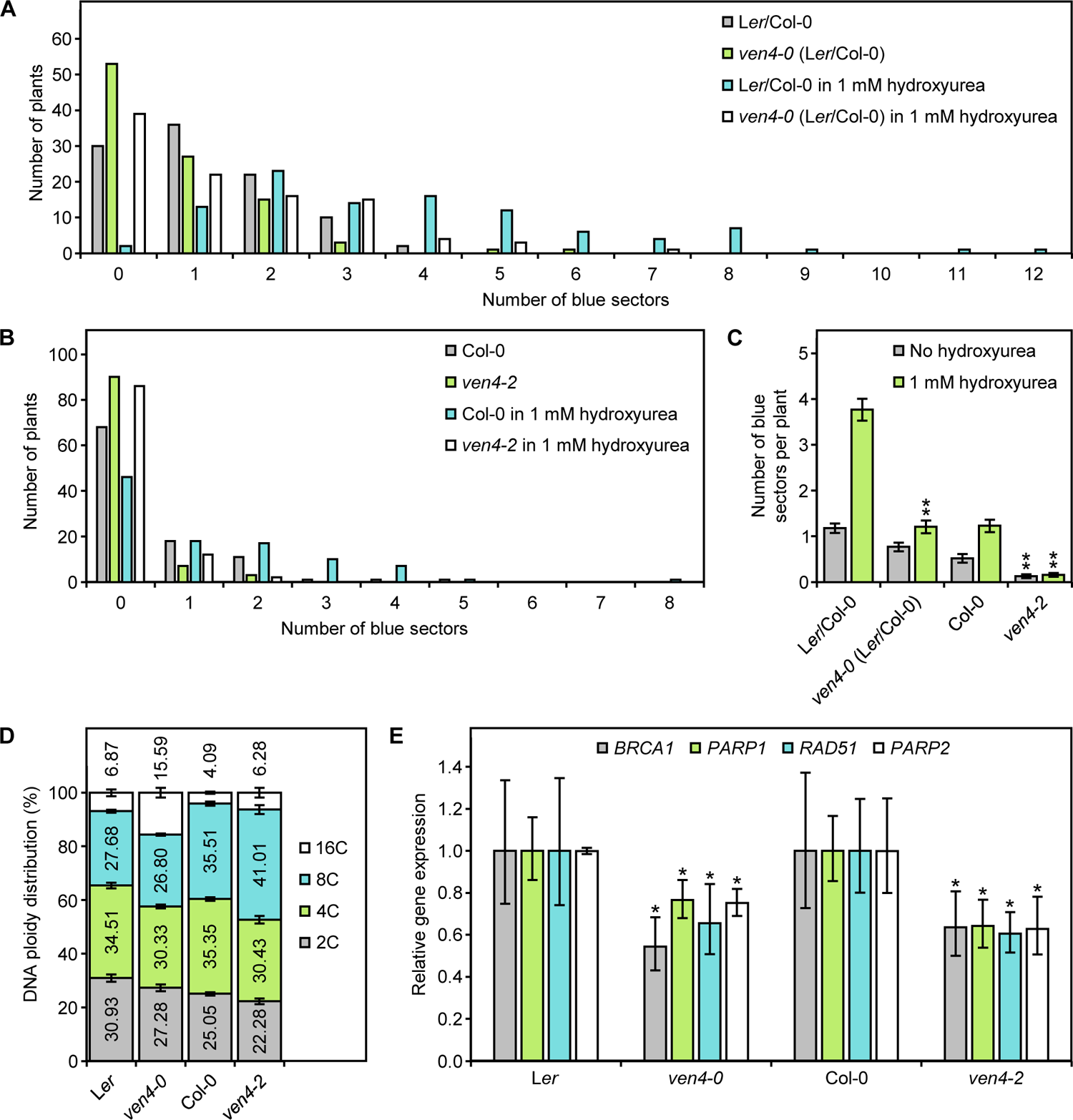
*ven4* mutants display signs of DNA damage and impaired DNA repair by HR. (A–C) Spontaneous and hydroxyurea-induced HR events in the *ven4-0* and *ven4-2* mutants. (A and B) Distribution of blue sectors visualized by GUS staining in *ven4* or phenotypically wild-type (*VEN4/VEN4* or *VEN4/ven4*) F3 plants derived from *ven4-0* (in a L*er* genetic background) × *IU.GUS 8* (Col-0) (A) and *ven4-2* (Col-0) × *IU.GUS 8* (Col-0) (B) crosses; plants were grown on medium not supplemented or supplemented with 1 mM hydroxyurea. Plants were collected at 12 das. (C) Mean number of blue sectors per F3 plant. Error bars indicate standard deviation. At least 100 plants from two different F3 families were analyzed. Asterisks indicate values significantly different from those of the corresponding phenotypically wild-type F3 plants in a Student’s *t*-test (\**P* < 0.001). (D) Nuclear ploidy distribution in third-node leaves of L*er*, *ven4-0*, Col-0, and *ven4-2* plants. Error bars indicate standard deviation. Three biological replicates of 100,000 events each were analyzed per genotype. Leaves were collected at 15 das. (E) RT-qPCR analysis of the relative expression of *BRCA1*, *PARP1*, *RAD51*, and *PARP2* in L*er*, *ven4-0*, Col-0, and *ven4-2*. Error bars indicate the interval delimited by 2^-ΔΔC^T ^± SD^, where SD is the standard deviation of the ΔΔCT values. Three biological replicates were examined per genotype, with three technical replicates in each experiment. The *ACT2* gene served as an internal control for normalization and relative quantification of gene expression. Asterisks indicate ΔCT values significantly different from those of the corresponding wild type in a Student’s *t*-test (\**P* < 0.05).

### Endoreduplication is increased and DNA repair genes are downregulated in *ven4* mutants

DNA damage has been associated with a reduced division rate in animal and plant cells because cell division halts until specific DNA repair mechanisms are activated (reviewed by Sancar et al.^58^ and Toettcher^59^). Cells from certain plants, such as Arabidopsis, use endoreduplication as a response to DNA damage, including DSBs^60^, a mechanism that is used less frequently by animal cells (reviewed by Fox and Duronio^61^ and Nisa et al.^62^). As DNA DSB repair by HR is impaired in *ven4* mutants, we examined their ploidy levels. We found lower 2C and 4C ploidy levels and higher 16C levels in *ven4-0* and *ven4-2* cells compared with L*er* and Col-0 cells, respectively (Figure 3D), resulting in an endoreduplication index^63, 64^ that was 19% and 10% greater than their wild types, respectively.

Expression of *TSO2*, encoding the major contributor to Arabidopsis RNR activity, is induced by bleomycin, a genotoxic agent that causes DNA DSBs, but not by hydroxyurea.^52, 65^ TSO2 is primarily nuclear in meristematic cells and cytoplasmic in adult cells. However, in quiescent cells, TSO2 is not detected under normal conditions but accumulates in the nucleus after exposure to ultraviolet (UV)-C radiation, suggesting a role for this protein in DNA damage protection and/or repair.^65^ Supporting this hypothesis, the *POLY (ADP-RIBOSE) POLYMERASE 1* (*PARP1*), *PARP2*, and *RADIATION SENSITIVE 51* (*RAD51*, which encodes a recombinase) genes, which play key roles in DNA DSB repair, are up-regulated in *tso2-1 rnr2a* double mutant plants.^54^ *BREAST CANCER SUSCEPTIBILITY1* (*BRCA1*), *RAD51*, *PARP1*, and *PARP2* are also up-regulated when DNA damage is induced and in several Arabidopsis mutants with impaired DNA repair activity, such as *photoperiod-independent early flowering1-3* (*pie1-3*), *actin-related protein6-3* (*arp6-3*), *SWR1 complex6-1* (*sw6-1*), *jing he sheng 1* (*jhs1*), or *retinoblastoma-related 1* (*rbr1*).^66–68^

As previously mentioned, we observed that *ven4* mutants are hypersensitive to hydroxyurea and show reduced ability to repair DSBs by HR. In addition, since human SAMHD1 is involved in the resolution of DSBs by both HR^14^ and NHEJ^69^, it is likely that VEN4 also has a role in DNA repair. To study whether DDR is altered in *ven4* mutants, we analyzed the expression of some of the above-mentioned key DNA repair genes by reverse transcription quantitative PCR (RT-qPCR). Interestingly, the mRNA levels of the DNA damage-induced genes *BRCA1*, *RAD51*, *PARP1*, and *PARP2* were significantly lower in *ven4-0* and *ven4-2* mutants than in their L*er* and Col-0 wild types when grown in the absence of genotoxic drugs, which suggests a direct or indirect role for VEN4 in the basal induction of these genes (Figure 3E).

### R-loops do not accumulate in *ven4* mutants

Recent work showed that human SAMHD1 functions as a tumor suppressor by resolving R-loops that often form at regions of the genome where transcription and replication conflict. Indeed, even in the absence of DNA damage inducers, R-loops accumulate in cells lacking SAMHD1 function, increasing replication stress and genome instability, and triggering DDRs.^16^

To determine whether this function of human SAMHD1 is conserved in VEN4, we performed a dot blot analysis using the well-established S9.6 monoclonal antibody^70^ to detect R-loops in total DNA extracted from *ven4-0* and *ven4-2* plants grown in the presence or absence of 1 mM hydroxyurea and collected at 19 das (Figure 2A–J). Given the previously described cross-reactivity of the S9.6 antibody with single-stranded (ss) or double-stranded (ds) RNAs^71^, we employed RNase T1 and RNase III to cleave ssRNAs and dsRNAs, respectively. Additionally, we used RNase H, which specifically cleaves RNAs within R-loops, thus ensuring the specificity of the S9.6 antibody signal. Simultaneously, we employed an anti-dsDNA antibody as a loading control. However, unlike in human cells lacking SAMHD1 function, there appeared to be no substantial accumulation of R-loops in the *ven4-0* and *ven4-2* mutants compared with their L*er* and Col-0 wild types, even when grown in the presence of the DNA-replication-blocking agent hydroxyurea (Figure S4).

### The *ven4-0* and *tso2-1* mutants have less fumarate and malate

Fumarate, malate, and succinate have been identified as competitive inhibitors of some α-ketoglutarate-dependent enzymes involved in DDRs across human, yeast, and bacterial cells.^72–76^ Fumarate has also been identified as a defense-priming agent against pathogens in Arabidopsis^77^, but its association with the DDR has not yet been established; nor has that of malate or succinate.

Interestingly, metabolite profiling analysis of the *ven4-0* mutant detected lower levels of fumarate and malate than in the L*er* wild type. The *tso2-1* mutant, carrying a loss-of-function allele of *TSO2*, also exhibited lower levels of fumarate, malate, and succinate than L*er* (Table S2). These findings support the hypothesis of a direct or indirect role for VEN4 in the DDR and additionally point to the conserved functions of fumarate, malate, and/or succinate in DNA repair in plants.

## DISCUSSION

### Arabidopsis VEN4 and human SAMHD1 might play similar roles not only in dNTP metabolism but also in maintenance of genome stability

Here, we examined VEN4, the likely Arabidopsis ortholog of human SAMHD1^27–29^ (Figures 1 and S1), which is highly conserved in plants, but lacks the SAM domain present in SAMHD1 (Figure S2). Since conservation of the dNTPase-dependent roles of SAMHD1 and VEN4 has already been stablished^27–29^, we aimed to determine whether VEN4 has also a dNTPase-independent role, participating in the maintenance of genome stability, particularly in DNA DSB repair by HR and resolution of R-loops, similar to human SAMHD1.^13–16, 35, 45^

DNA DSBs are considered the most harmful genetic lesions in animal and yeast cells; they can be induced by exposure to exogenous genotoxic agents or chemicals, such as ionizing radiation or hydroxyurea, but can also arise spontaneously due to cellular metabolism (reviewed by Chapman et al.^78^). In animals, DDRs ensure genome integrity by arresting cell cycle progression to aid DNA repair and, when necessary, by inducing apoptosis. In vascular plants, endoreduplication enables cells to duplicate their genomes without division, thereby increasing ploidy levels. This mechanism effectively prevents the transmission of damaged DNA to daughter cells.^79^ For instance, in Arabidopsis, endoreduplication is activated in response to DNA DSBs, but not in direct response to replication stress.^60^ Indeed, in the absence of DNA damage inducers, we found that *ven4* mutants exhibit increased ploidy levels in leaf cells, suggesting an over-accumulation of spontaneous DSBs.

In eukaryotes, transcription of genes encoding RNR subunits is activated in the S phase of the cell cycle and in response to DNA damage (reviewed by Guarino et al.^80^). Furthermore, to overcome the accumulation of DNA lesions, DDR genes are induced under genotoxic conditions.^52^ Using a *VEN4pro:GUS* transgene, we detected transcriptional induction of *VEN4* upon hydroxyurea treatment, supporting the hypothesis that VEN4 acts in DNA repair. Interestingly, the *ven4* mutants exhibited repression of key genes involved in DSB repair (*BRCA1*, *RAD51*, *PARP1*, and *PARP2*) when grown in culture medium not supplemented with hydroxyurea, suggesting a direct or indirect role for VEN4 in the basal transcriptional induction of these genes. Indeed, *IU.GUS* transgenic lines in the *ven4* mutant background showed impaired HR when grown in the absence of genotoxic drugs, which worsened following supplementation with 1 mM hydroxyurea, pointing to a role for VEN4 in spontaneous or induced DSB repair by HR, similar to its human ortholog.

Comparing the amino acid sequences and domain structures, along with the understanding of SAMHD1 function, can help us infer how VEN4 functions, and which of these functions are conserved with SAMHD1. Deacetylation of the K354 residue of human SAMHD1, located in the middle of its HD domain, occurs in response to DSBs. Deacetylation of K354 facilitates SAMHD1 binding to ssDNA at DSBs, which also facilitates CtIP binding to ssDNA in order to promote DNA end resection and HR.^35^ Interestingly, the *ven4-0* mutation causes a E249L substitution in the residue adjacent to the K248 of VEN4, which is equivalent to the K354 of human SAMHD1 and is highly conserved in other plants. *In silico* predictions of the effects of the E249L substitution on VEN4 and of the equivalent E355L mutation on human SAMHD1, revealed an increase in conformational stability and a decrease in flexibility^27^, which might impair the deacetylation of K248 (K354 in human SAMHD1) and the ability of the mutant proteins to bind ssDNA at DSBs. This hypothesis is consistent with the over-accumulation of both spontaneous and induced DSBs suggested by the impaired HR revealed by our *IU.GUS* assays, as well as by the repression of genes involved in DNA DSB repair.

### Fumarate and malate depletion in the *ven4-0* mutant also supports the implication of VEN4 in the DDR

Fumarase is a highly conserved enzyme best known for catalyzing the hydration of fumarate to L-malate in the tricarboxylic acid (TCA) or Krebs cycle in mitochondria, facilitating a transition step in the production of energy for the cell. However, a cytoplasmic echoform^81^ of this enzyme has been found in eukaryotes. Initially proposed to function as a scavenger of fumarate from the urea cycle and amino acid catabolism, this fumarase echoform has also been associated with the cellular DDR to DSBs (reviewed by Leshets et al.^82^). This role requires the movement of cytoplasmic fumarase to the nucleus in response to DNA damage, where it catalyzes the reverse conversion of L-malate to fumarate, leading to a local accumulation of fumarate at DSBs, which promotes their repair via HR or NHEJ by inhibiting some α-ketoglutarate-dependent histone demethylases.^75, 76^

Indeed, yeast and human cells lacking fumarase are hypersensitive to some DNA DSB and replication-block inducers, such as ionizing radiation or hydroxyurea.^74^ The fumarase of the prokaryote *Bacillus subtilis* also appears to participate in the DDR, achieved in this case by the biosynthesis of L-malate, which affects the translation and subcellular localization of the first protein to be recruited to DSB sites.^73^ Similar to fumarate, succinate appears to play a role in the control of epigenetic marks following DNA damage in yeast and human cells, but also in *Escherichia coli*.^72, 75^ By acting as structural mimics to α-ketoglutarate, fumarate and succinate competitively inhibit certain α-ketoglutarate-dependent enzymes involved in the successful repair of damaged DNA. Consequently, they have been defined as oncometabolites (reviewed by Liu and Yang^83^).

The *ven4-0* point mutation has been particularly useful for predicting the effects of VEN function in DNA DSB repair, strongly reinforcing our other experimental results. In particular, we found a depletion of fumarate and malate in this mutant, which may be partly related to the repression of key DNA repair genes when plants are grown in medium not supplemented with hydroxyurea, as previously described in *E. coli*, where the loss of function of fumarase affects transcriptional induction of many DNA-damage-responsive genes.^72^

### VEN4 does not appear to prevent R-loop accumulation

Human SAMHD1 resolves R-loops at genomic regions where the transcription and replication machineries collide.^15, 16^ This role of SAMHD1 is independent of its dNTPase activity. Our results support the hypothesis of a conserved role for VEN4 in maintaining genome integrity, specifically in DNA DSB repair by HR. However, we failed to obtain evidence that VEN4 resolves R-loops, even upon genotoxic treatment with hydroxyurea. This may be the correct result, or the dot blots may only have enough sensitivity to detect major changes in R-loop accumulation and not small and/or localized changes in R-loop formation at transcription–replication conflict regions of the genome. Thus, despite our negative results, we cannot definitively conclude that VEN4 does not participate, directly or indirectly, in the resolution of R-loops at these conflictive genomic regions, as SAMHD1 does in humans.

### Limitations of the study

Therefore, in summary, we show that, like SAMHD1, Arabidopsis VEN4 functions in DNA repair. However, whether it functions in resolving R-loops remains to be resolved by additional studies. Examining how the differing domain structures of the two proteins (VEN4 lacks the SAM domain present in SAMHD1 but contains other conserved domains) affect their respective functions also remains an intriguing question for future research. Finally, it will be interesting to decipher the mechanism by which these proteins function in DNA repair, and how the lack of VEN4 affects plant development. In addition to opening these new avenues of inquiry, our study provided insight on the conservation of DNA repair mechanisms and their relationship to dNTP metabolism, in eukaryotes.

## ACKNOWLEDGEMENTS

The authors wish to thank J.M. Serrano, M.J. Ñíguez, J. Castelló, C. Yamaguchi, and R. Sasaki for their excellent technical assistance, and Ortrun Mittelsten Scheid and Zhongchi Liu for providing seeds.

## AUTHOR CONTRIBUTIONS

M.R.P. and J.L.M. conceived and supervised the study, provided resources, and obtained funding. R.S.-M., H.T., V.Q., J.L.M., and M.R.P. designed the methodology. R.S.-M., S.F.-C., K.K., and A.O. performed the research. R.S.-M., S.F.-C., J.L.M., and M.R.P. wrote the original draft. All authors reviewed and edited the manuscript. J.L.M. and M.R.P. contributed equally to this work.

## METHODS

### RESOURCE AVAILABILITY

#### Lead contact

Further information and requests for resources and reagents should be directed to and will be fulfilled by the lead contact, María Rosa Ponce.

#### Materials availability

Materials generated in this study will be made available on request. For further details contact the lead contact.

#### Data and code availability

Sequence data from this article can be found at TAIR (http://www.arabidopsis.org) under the following accession numbers: *VEN4* (At5g40270), *TSO2* (At3g27060), *BRCA1* (At4g21070), *RAD51* (At5g20850), *PARP1* (At2g31320), *PARP2* (At4g02390), and *ACT2* (At3g18780). This paper does not report original code. Any additional information required to reanalyze the data reported in this paper is available from the lead contact upon request.

## EXPERIMENTAL MODEL AND STUDY PARTICIPANT DETAILS

The *Arabidopsis thaliana* (L.) Heynh. wild-type accessions Col-0 and L*er*, and the T-DNA insertional mutants *ven4-2* (SALK_077401) and *ven4-3* (SALK_131986), in the Col-0 genetic background, were originally obtained from the Nottingham Arabidopsis Stock Centre (NASC; Nottingham, United Kingdom) and then propagated in the laboratory for further analysis. The EMS-induced *ven4* mutant, in the L*er* genetic background, was isolated in the laboratory of José Luis Micol^23, 84–86^ and renamed recently as *ven4-0* by Sarmiento-Mañús *et al.*^27^ Seeds of the *IU.GUS 8* transgenic line^57^, in the Col-0 genetic background, were kindly provided by Ortrun Mittelsten Scheid (Gregor Mendel Institute of Molecular Plant Biology, Austrian Academy of Sciences, Vienna BioCenter, Vienna, Austria), and *tso2-1*^54^ seeds, in the L*er* genetic background, by Zhongchi Liu (University of Maryland, College Park, MD, USA).

## METHOD DETAILS

### Plant growth conditions and genotyping

Except for metabolite profiling (see below), plants were grown under sterile conditions on half-strength Murashige and Skoog (MS; Duchefa Biochemie) medium containing 1% (w/v) sucrose (Duchefa Biochemie), 0.1% (w/v) 2-Morpholinoethanesulfonic acid (MES) monohydrate (Duchefa Biochemie), and 0.7% (w/v) plant agar (Duchefa Biochemie) at 20°C ± 1°C, 60–70% relative humidity, and continuous fluorescent light of ∼75 μmol·m^−2^·s^−1^, as previously described by Ponce et al.^87^ and Berná et al.^23^ When required, MS medium was supplemented with 15 µg·mL^−1^ of hygromycin B (Invitrogen). Mutants were genotyped by PCR amplification and/or Sanger sequencing, as previously described by Sarmiento-Mañús *et al.*^27^, using the primers shown in Table S1. Unless specified otherwise, all plants employed in this work were homozygous for the indicated mutations.

### Protein structure visualization and bioinformatic analyses

The 3D structures of the Arabidopsis VEN4 and human SAMHD1 proteins shown in Figure 1 were obtained from the AlphaFold Protein Structure Database (AlphaFold DB; Jumper et al.^88^; Varadi et al.^89^; https://alphafold.ebi.ac.uk/), where they are identified as AF-Q9FL05-F1 and AF-Q9Y3Z3-F1, respectively, and visualized using UCSF ChimeraX 1.2.5 (Goddard et al.^90^; Pettersen et al.^91^; https://www.rbvi.ucsf.edu/chimerax/). Root mean square deviations (RMSD) for both proteins were determined with the MatchMaker function of UCSF ChimeraX 1.2.5, using default parameters. The multiple sequence alignments shown in Figures S1 and S2 were obtained using ClustalX 1.5b.^92, 93^ The identical and similar residues among the analyzed protein sequences were shaded in black and gray, respectively, using BoxShade (https://junli.netlify.app/apps/boxshade/).

### Hydroxyurea treatment

The hydroxyurea hypersensitivity of the *ven4* mutants was measured as the percentage of primary root growth inhibition. Plants were grown for 19 das on vertically oriented Petri dishes with half-strength MS medium supplemented or not with 1 mM hydroxyurea. Primary root length was measured from photographs taken of plants grown on those Petri dishes using the NIS Elements AR 3.1 image analysis software (Nikon). To analyze rosette morphological phenotypes, plants were grown for 19 das on horizontally oriented Petri dishes with half-strength MS medium supplemented or not with 1 mM hydroxyurea.

### GUS staining and analysis of *IU.GUS* transgenic plants

Plants homozygous for the *VEN4pro:GUS* transgene in a Col-0 genetic background^27^ were collected at 11 das from two different T2 families grown on half-strength MS medium supplemented or not with 1 mM hydroxyurea. For DNA repair analyses, Col-0, *ven4-0*, and *ven4-2* plants were crossed with the *IU.GUS 8* transgenic line^57^, and 100 Hyg^R^ seedlings (50 phenotypically mutant and 50 phenotypically wild type) from two different F3 families derived from each cross were collected at 12 das. GUS enzymatic activity in *VEN4pro:GUS* and *IU.GUS* transgenic plants was visualized as previously described by Robles et al.^94^ Blue sectors in *IU.GUS* transgenic plants were counted using a MZ6 stereomicroscope (Leica) equipped with a DXM1200 digital camera (Nikon).

### Estimation of chloroplast-to-nuclear genome copy ratio

Chloroplast-to-nuclear genome copy ratios were estimated by qPCR using genomic DNA from L*er*, Col-0, *ven4-0*, and *ven4-2* plants at 15 das and the At4g04930_F/R and ArthCp025_F/R primer pairs (Table S1) for amplification of the nuclear At4g04930 and chloroplast ArthCp025 single-copy genes, respectively, as first described by Garton *et al.*^47^ The relative ratios of chloroplast-to-nuclear genome copies were subsequently calculated according to Yoo *et al.*^49^, normalizing to the wild-type values, which were set to 1.

### Flow cytometry

Nuclei were extracted from plants collected 7 das as previously described by Horiguchi et al.^95^ Briefly, first- and second-node leaves were chopped with a razor blade in 500 μL of cold (kept on ice) Galbraith’s nuclear extraction buffer.^96^ The cell suspension was then filtered through a 30 μm pore size nylon mesh, treated with 100 μg·mL^−1^ of RNase A (Roche), and stained with 50 μg·mL^−1^ of propidium iodide (PI; Sigma-Aldrich) for 10 min. Nuclear DNA content was measured by flow cytometry using a SH800S Cell Sorter (Sony). Three biological replicates of 100,000 events each were analyzed per genotype, and the endoreduplication index (*EI*) was calculated as *EI* = (0 × %2C) + (1 × %4C) + (2 × %8C) + (3 × %16C) + (4 × %32C), as previously described by Sterken *et al.*^63^

### Total RNA isolation and RT-qPCR experiments

For gene expression analysis, total RNA from plant rosettes harvested at 10 das was extracted with TRIzol (Invitrogen). Reverse transcription and qPCR amplification were performed as described by Wilson-Sánchez et al.^97^ qPCRs were carried out in a Step-One Real-Time PCR System (Applied Biosystems) using three technical replicates per biological replicate (each consisting of three plant rosettes). The primers used are listed in Table S1. The housekeeping gene *ACTIN2* (*ACT2*) served as an internal control for normalization and relative quantification of gene expression. The CT values were normalized using the 2^−ΔΔCT^ method.^98^

### R-loop recognition by the S9.6 monoclonal antibody

Genomic DNA from ∼200 mg of plant rosettes harvested at 19 das was extracted using a DNeasy Plant Mini Kit (Qiagen) according to the manufacturer’s instructions, except that use of the RNase A included in the kit was omitted. The isolated DNA was quantified using a Qubit 2.0 Fluorometer and a Qubit dsDNA High Sensitivity (HS) Assay Kit (Invitrogen). Samples of 1.2 µg of DNA were treated with either 5 U of RNase H (New England Biolabs), 1,000 U of RNase T1 (Thermo Scientific), or 1 U of Ambion RNase III (Invitrogen) for 2 h at 37°C to assess the specificity of the anti-DNA–RNA hybrid (S9.6) antibody (AB_2861387; Sigma-Aldrich). An untreated sample was also included. Samples of 25, 50, or 100 ng of DNA (in a volume of 2 µL), either treated or untreated with RNases, were blotted onto two positively charged nylon membranes, one for the S9.6 antibody and the other for the anti-dsDNA antibody (AB_470907; Abcam). The samples were allowed to saturate into the membranes before crosslinking with UV light (1,200 μJ × 100). The membranes were blocked with 5% (w/v) skimmed milk in 1× Tris-Buffered Saline solution with 0.05% (v/v) Tween-20 (1× TBST) for 1 h at room temperature on a shaker. After three subsequent washes with 1x TBST, the membranes were incubated overnight at 4°C on a shaker with either the S9.6 or the anti-dsDNA antibody, at 1:1,000 or 1:10,000 dilution, respectively, in 1× TBST. Following three washes with 1x TBST, the membranes were incubated with WesternSure goat anti-mouse IgG HRP-conjugated secondary antibody (AB_2721263; LI-COR), diluted at 1:50,000 in 1× TBST, for 2 h at room temperature on a shaker. After rinsing the membranes in 1× TBST, antibodies were detected using WesternSure PREMIUM Chemiluminescent Substrate (LI-COR) and a C-Digit Blot Scanner (LI-COR).

### Metabolite profiling

The first leaves of L*er*, *ven4-0*, and *tso2-1* plants grown on rock wool at 22°C under a long-day photoperiod (16-h light/8-h dark) were collected at 14 das from three biological replicates. After harvesting, 20–80 mg of tissue was immediately frozen in liquid nitrogen and stored at −80°C until further use. Metabolite profiling was performed by capillary electrophoresis time-of-flight mass spectrometry (CE-TOF MS), as described by Oikawa et al.^99, 100^ and Ferjani et al.^101^ The raw CE-TOF MS data were processed, and metabolite peaks were identified, aligned, and annotated using MasterHands.^102^ Metabolites with significantly different levels between L*er* and *ven4-0* or *tso2-1* plants were identified using a Student’s *t*-test (Table S2).

## QUANTIFICATION AND STATISTICAL ANALYSIS

Quantification and statistical analyses are described in the Method Details and Results sections as well as in the figure legends.

## SUPPLEMENTAL INFORMATION

**Figure S1.** Amino acids and the HD motif are conserved in the VEN4 protein.

**Figure S2.** Key residues are conserved in putative plant orthologs of the Arabidopsis VEN4 protein.

**Figure S3.** Hydroxyurea affects root growth of *ven4* mutants.

**Figure S4.** R-loops do not accumulate in *ven4* mutants.

**Table S1.** Primers used in this work.

**Table S2.** Metabolite profiling results.

