## Supplementary material for "Functional conservation and divergence of Arabidopsis VENOSA4 and human SAMHD1 in DNA repair": Figures S1-S4 and Table S1

### Supplemental Figures and Tables

Supplemental Material included in this file:

- Figures S1-S4
- Table S1

Supplemental Material not included in this file:

- Table S2

```
Homo sapiens              1 MQRADSEQPSKRPRCDDSPRTPSNTPSAEADWSPGLELHPDYKTWGPEQVCSFLRRGGFE
Arabidopsis thaliana      1 ------------------------------------------------------------

Homo sapiens             61 EPVLLKNIRENEITGALLPCLDESRFENLGVSSLGERKKLLSYIQRLVQIHVDTMKVIND
Arabidopsis thaliana      1 -----------------MGAYCDENLSSLPVFSSGAP-----------ANELRFSKHVYD

Homo sapiens            121 PIHGHIELHPLLVRIIDTPQFQRLRYIKQLGGGYYVFPGASHNRFEHSLGVGYLAGCLVH
Arabidopsis thaliana     33 NVHGNIYLDPLCLKFIDTEQFQRLRELKQLGVTNMVYPGAVHSRFEHSLGVYWLAGETAQ

Homo sapiens            181 ALGEK-QPELQISERDVLCVQIAGLCHDLGHGPFSHMFDGRFIPLARPEVKWTHEQGSVM
Arabidopsis thaliana     93 RLKNFQGMELGIDNYDLQTVRLAGLLHDIGHGPFSHMFEREFLPKVISDCQWSHELMSVN

Homo sapiens            240 MFEHLINSNGIKPVMEQYGLIPEEDICFIKEQIVGPLESPVEDSLWPYKGRPENKSFLYE
Arabidopsis thaliana    153 MIDHMVDTHHIDIDAQMLKR--------VKDMIL-------ASTEFSQLKGNAEKRFLYD
                              ven4-2

Homo sapiens            300 IVSNKRNGIDVDKWDYFARDCHHLGIQNNFDYKRFIKFARVCEVDNELRICARDKEVGNL
Arabidopsis thaliana    198 IVANGRNGIDVDKFDYLVRDSRACGLGSNFQFQRLTETMRVLD----NEICFRAKEYLSV
                                                                                ven4-0

Homo sapiens            360 YDMFHTRNSLHRRAYQHKVGNIIDTMITDAFLKADDYIEITGAGGKKYRISTAIDDMEAY
Arabidopsis thaliana    254 HKLFATRADLYRTVYTHSKVKAIELMIVDAMVKANNHLE----------ISSMINDPSEY

Homo sapiens            420 TKLTDNIFLEILYSTDPKLKDAREILKQIEYRNLFKYVGETQPTGQIKIKREDYES-LPK
Arabidopsis thaliana    304 WKLDDTILKTIEIAPDPELAEAKELILRVRRRQLYQFCNEYAVPKD---KIDHFKAVTPQ

Homo sapiens            479 EVASAKPKVLLDVKLKAEDFIVDVINMDYGMQEKNPIDHVSFYCKTAPNRAIRITKNQVS
Arabidopsis thaliana    361 DIICS--QKHTSLTLKEEDIAVTNVKIDLARGRENPLECINFYKDYDSAEKFVIPEDRVS
                                                                              ven4-3

Homo sapiens            539 QLLPEKFAEQLIRVYCKKVDRKSLYAARQYFVQWCADRNFTKPQDGDVIAPLITPQKKEW
Arabidopsis thaliana    419 HLLPTTYQDMIVRVYAKKPELVEAVSE--AFEN------FQMRTYGIKAQVHATPEKKKR

Homo sapiens            599 NDSTSVQNPTRLREASKSRVQLFKDDPM
Arabidopsis thaliana    471 RVM-------------------------
```

**Figure S1.** Amino acids and the HD motif are conserved in the VEN4 protein. Related to Figure 1. Residues conserved between the human SAMHD1 and Arabidopsis VEN4 sequences are shaded in black, while similar residues are shaded in gray. Numbers indicate amino acid positions. Triangles, red letters, and pink and blue underlines indicate the *ven4-2* and *ven4-3* T-DNA insertions, the conserved E residue altered by the *ven4-0* point mutation, and the human SAM and HD domains, respectively. Orange, blue, green, pink, and purple letters represent conserved residues with a functional role in SAMHD1 described previously (R333, salt bridge with E355 and SAMHD1 dNTPase activity [Ji et al., 2013, 2014; Sarmiento-Mañús et al., 2023]; K354, DNA DSB repair by HR [Kapoor-Vazirani et al., 2022]; R451/L453, RXL motif, cyclin A2, CDK1, or CDK2 interactions [St Gelais et al., 2018]; K484, CtIP interaction for DNA end resection by HR at DSB sites [Cabello-Lobato et al., 2017; Daddacha et al., 2017]; T592, stalled DNA replication forks resection and R-loops resolution [Cribier et al., 2013; White et al., 2013; Coquel et al, 2018]).

```
Medicago truncatula         ------------------------------------------------------------
Olea europaea               ------------------------------------------------------------
Arabidopsis thaliana        ------------------------------------------------------------
Oryza sativa             1  MKHPSRIKLASPNKSHGLPYLLPVSAGFLPFSHRLACLLAFPIPISHQPQTAANPTSARP
Quercus lobata              ------------------------------------------------------------
Solanum lycopersicum        ------------------------------------------------------------

Medicago truncatula      1  -----------MGAYHNDAVSLP-------PRYDVVSSRD--------------------
Olea europaea            1  -----------MGAFCNDDIAFSNFNFG--SFQDLRSS----------------------
Arabidopsis thaliana     1  -----------MGAYCDENLSSLPVFSSGAPANELRFS----------------------
Oryza sativa            61  HRNLARFSEEEMGEYCGAAPEEDPAMALVTPLPTTTTTTTTAAAAAIKQPHYYGCFDRCS
Quercus lobata           1  -----------MGLSTNFEDSTD-----------RRLL----------------------
Solanum lycopersicum     1  -----------MGDCSNSKLLPMCD-SGEAPFDTRRYY----------------------

Medicago truncatula     23  -KHVHDNVHGNIFIDSLSLKFIDTEQFQRLRELKQLGFTHLVYPGAVHSRFEHSLGVYWI
Olea europaea           26  -KHVYDNVHGNIYLGPLFLKFIDTEQFQRLRDLKQLGMAHMVYPGAVHSRYEHSLGVYWL
Arabidopsis thaliana    28  -KHVYDNVHGNIYLDPLCLKFIDTEQFQRLRELKQLGVTNMVYPGAVHSRFEHSLGVYWL
Oryza sativa           121  TKQVFDNLHGNISLDPLAREFVDTEEFQRLRDLKQLGLTYLVYPGAVHTRFEHSLGVYWL
Quercus lobata          17  -KQVHDNVHGNIYLEPVALKFIDTEEFQRLRELKQLGLSNMVYPGAVHSRFEHSLGVYWL
Solanum lycopersicum    27  -KHIHDNVHGNIYLDKQALNFIDTEQFQRLRELKQLGLGYMVYPGAVHSRFEHSLGVYWL

Medicago truncatula     82  AGQSVEKLNSYQGMELGIDKFDIQSVKLAGLLHDVGHGPFSHLFEREFLPRVISGSHWSH
Olea europaea           85  AGEAVHKLKNSHGLEHGIDHFDVQTVKLAGLLHDVGHGPFSHLFEREFLPMVHKGSEWSH
Arabidopsis thaliana    87  AGETAQRLKNFQGMELGIDNYDLQTVRLAGLLHDIGHGPFSHMFEREFLPKVISDCQWSH
Oryza sativa           181  AGEAMNNLRLYQGEELGIDRVDMQTVKLAGLLHDIGHGPFSHLFEHEFLPRVVPGSTWTH
Quercus lobata          76  AGKAVDTIKTCQGLELGIERSDIKLVKLAGLLHDVGHGPFSHMFEKEFLPRVFNGFPWSH
Solanum lycopersicum    86  ASDAVHRLKTYQGQELGIESFDVQTVKLAGLLHDVGHGPFSHLFEREILPRVRSGIKWSH

Medicago truncatula    142  EQMSVKMVDYIVGEHHIDIDPHMLKRVKEMILASSEFSLPRSSSEKGFLYDIVANGRNGI
Olea europaea          145  EQMSVDMVDHIVDEHHIEIESEALKRVKEMILASSKYATTKSTREKHFLYDIVANGRNGI
Arabidopsis thaliana   147  ELMSVNMIDHMVDTHHIDIDAQMLKRVKDMILASTEFSQLKGNAEKRFLYDIVANGRNGI
Oryza sativa           241  ENMSALLLDSIVDKHQIDIEADHLKIVMEMIVASSKFTATESTKEKRFLYDIVANGRNGI
Quercus lobata         136  EDMSVKMVDHIVDEHNIDIDSESLTKVKQMITASSEHDSTENMKEKKFLYDIVANGRNGI
Solanum lycopersicum   146  EDMSLKMIDYIVDENSIDIDSGTLKKVKEMIVASEAGKSVSS-KEKQFLYDIVANGRNGI

Medicago truncatula    202  DVDKFDYIARDCRACGLGCNFEFQRIMETMRVLDDEICYRAKDYLTIHKLFATRADLYRT
Olea europaea          205  DVDKFDYIVRDSRACGLGCNFQFQRIMETMRVMGDEICYRAKDYLTIHKLFATRADLYRT
Arabidopsis thaliana   207  DVDKFDYLVRDSRACGLGSNFQFQRLTETMRVLDNEICFRAKEYLSVHKLFATRADLYRT
Oryza sativa           301  DVDKFDYIGRDCRACGLGCNFQYWRLLQGMRVMGDEICYPAKDYLSIHKLFTTRADLHRT
Quercus lobata         196  DVDKFDYIVRDSRACGLGCNFHFERIMESMRVMGDEICYRAKDYLTIHKLFATRADLHRT
Solanum lycopersicum   205  DVDKFDYIERDTRACGLRCNFQFQRIMETMRVIDNEICYRAKEYLTIHKLFSTRADLHRT

Medicago truncatula    262  VYTHPKVKAMELMVVDALVQANDYLQISSSIQDPGEYWKLDDSIIKTIETSPLPELKEAR
Olea europaea          265  VYTHAKVKAIEFMVVDALLNANNYLEIASHIYDPSQYWKLDDTIIKTIETAPDHELKESR
Arabidopsis thaliana   267  VYTHSKVKAIELMIVDAMVKANNHLEISSMINDPSEYWKLDDTILKTIEIAPDPELAEAK
Oryza sativa           361  VYTHAKVKAVELMLVDALVEANEYLGIALHAQDPADFWKLDDTIIKSIETAPNDELNKAK
Quercus lobata         256  VYTHAKVKAIELMVVDALLKANDFLQIASSIRQPAEFWKLDDSILKIIEFSNAQELKEAR
Solanum lycopersicum   265  VYTHPKVKAIEFMVVDALIKANDHLEIDSYIDEPAKYWMLDDTIVKTIEASTHQDLEESR

Medicago truncatula    322  ELILRIRRRDLYQFCNEYAVPKDIMDNFKKVTPQDIVCSQKNGGVMLKEEDVAVCNVKID
Olea europaea          325  DLILRIRRRDLYQFCNEYAVPKDQLDNFKDVTAQDIVCSQKSGGVALKEEDVAVSNVRID
Arabidopsis thaliana   327  ELILRVRRRQLYQFCNEYAVPKDKIDHFKAVTPQDIICSQKHTSLTLKEEDIAVTNVKID
Oryza sativa           421  GIIQRIRRRELYKFCNEYSVPKDKLEHFKNITAQDIVCSQKSSKVLLKEEDVAVSNVKID
Quercus lobata         316  DIIQRIRRRELYQFCNEFSVPKDKMEHFKKITPQDIICSQKTGGVTLEEEDIVVSNVKID
Solanum lycopersicum   325  NLIRRIRRRDIYQYCNEFTVSKENLEYFKNVTAQDIICSQ-NSDAHLNEEDVIVTNIKID

Medicago truncatula    382  LTRGKHNPLESIHFFKDYESDEKFTIPDERISHLLPASFQDMIVRVYSKKPELVEKISEA
Olea europaea          385  LTRGKHNPLESIKFFKDYESDEKLSITDDRISHLLPTSYQDMIVRVYSKKPELVGAISEA
Arabidopsis thaliana   387  LARGRENPLECINFYKDYDSAEKFVIPEDRVSHLLPTTYQDMIVRVYAKKPELVEAVSEA
Oryza sativa           481  LTRGKDNPLESIKFFKDFGCDEKFPITDERVSHLLPAYNQDRIVRVYAKKHELVEAVSEA
Quercus lobata         376  LTRGRNNPLQRIMTAMRCSQSK--IASATCCLHLTKI-----------------------
Solanum lycopersicum   384  LANGRNNPLERVSFFQDYDSFEKFHIKEDCVSQLLPTCYQDLIVRVYARDPKLVDAVTNA

Medicago truncatula    442  FENYQLKTYGIKAQVHSTPDKKK-RRYNS-
Olea europaea          445  FENFQLKTYGIKAQVHSTPDKKK-RRIC--
Arabidopsis thaliana   447  FENFQMRTYGIKAQVHATPEKKK-RRVM--
Oryza sativa           541  FENLQLRMYGEKTQVHDTPRKKR-IRFH--
Quercus lobata              ------------------------------
Solanum lycopersicum   444  FENFQTKTYGEKTQVHAITEKKRRLKYNGN
```

**Figure S2.** Key residues are conserved in putative plant orthologs of the Arabidopsis VEN4 protein. Related to Figures 1 and S1. *Medicago truncatula* (XP_013450811.1); *Olea europaea* (CAA2994598.1); *Arabidopsis thaliana* (NP_568580.1); *Oryza sativa* (XP_015621837.1); *Quercus lobata* (XP_030961156.1); *Solanum lycopersicum* (XP_010320576.1). Amino acids of VEN4 equivalent to those of the HD domain of human SAMHD1 are underlined in blue. See the legend of Figure S1 for further details.

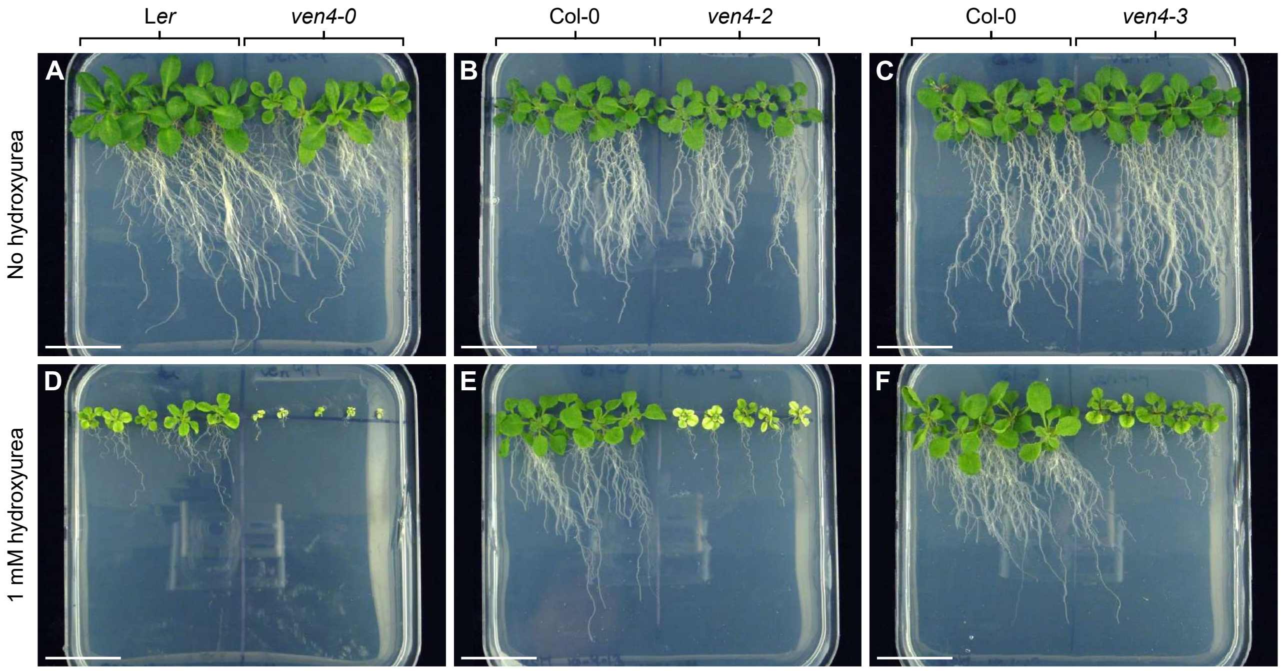


**Figure S3.** Hydroxyurea affects root growth of *ven4* mutants. Related to Figure 2. Plants were grown on medium not supplemented (A–C) or supplemented with 1 mM hydroxyurea (D–F). Photographs were taken 19 das. Scale bars: 3 cm.

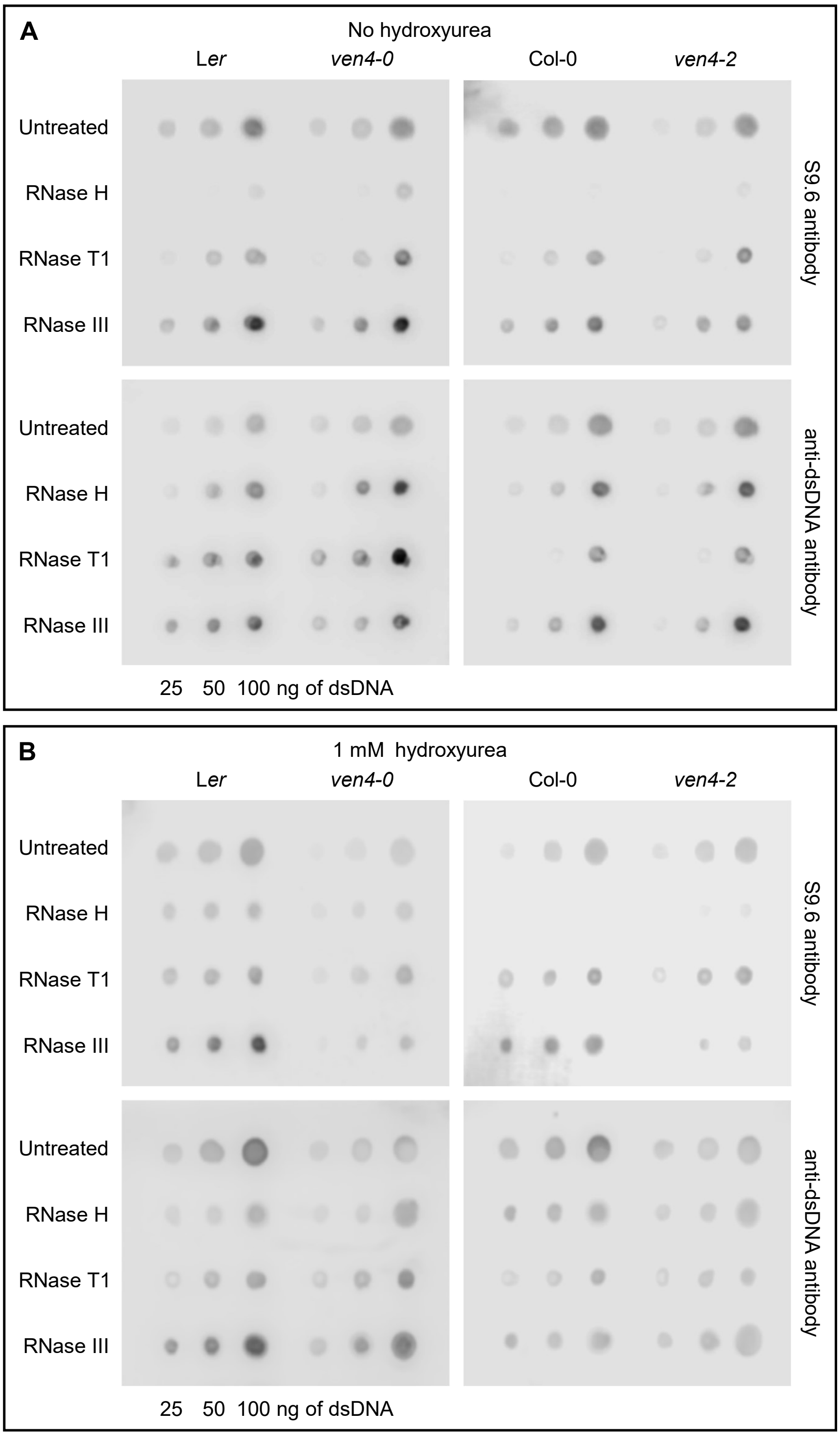


**Figure S4.** R-loops do not accumulate in *ven4* mutants. Related to Figures 2, 3, and S3. Global R-loops were detected by dot blot analysis with the S9.6 antibody in samples of 25, 50, and 100 ng of genomic DNA extracted from plants collected at 19 das that were grown on medium not supplemented (A) or supplemented (B) with 1 mM hydroxyurea. DNA samples were treated with either RNase H, RNase T1, or RNase III to assess the specificity of the S9.6 antibody before being blotted onto positively charged nylon membranes. An untreated DNA sample was also included. Simultaneous detection of dsDNA with an anti-dsDNA antibody was used as a loading control. This experiment was performed once.

**Table S1.** Primers used in this work. Related to STAR Methods.

| Purpose | | Oligonucleotide names | Oligonucleotide sequences (5′ → 3′) | |
|---|---|---|---|---|
| | | | Forward primer (F) | Reverse primer (R) |
| Genotyping of | *ven4-0* | At5g40270_F4/R4[a] | TTGATGTGGACAAGTTTGACTA | TGATTGTTGGCTTTGACCATGG |
| | *ven4-2* | At5g40270_F2/R2[a] | CTTAACTTTTAGTAGTTGGCTTCT | GTCTTCTGAGGAAGAACTGTAC |
| | *ven4-3* | At5g40270_F3/R3[a] | GTCTCGTTCCATTCATTTGCAG | CATGCTATATGTACACGCATCC |
| | *tso2-1* | TSO2_F/R[a] | CCTTCAATGCCAGAAGAGCC | TTCTTCAGCCAGAAGATTGAAC |
| T-DNA insertion verification | | LBb1.3[b] | ATTTTGCCGATTTCGGAAC | |
| qPCR | | At4g04930_F/R[c] | CAACCTTGCCTTTTCAACTC | CAATACCATCTACTCCTTGG |
| | | ArthCp025_F/R[c] | GAATTCCATCCGTTTTCTGG | AAGGCCACCCTATCCAACTC |
| RT-qPCR | | PARP1_F/R[d] | GTGTAGCGAAAAGATCTTGAAAGGAGAGG | CCATCCAGACAAACTTTCCAGTTCAGTAG |
| | | PARP2_F/R[d] | ACATGGTTTACACCAGATGGGGAAGAG | GGACTTGGGATGTGGGATAAACTCCTT |
| | | RAD51_F/R[d] | CTCCGAGGAAGGATCTCTTGCAG | GCTCGCACTAGTGAACCCCAG AGG |
| | | BRCA1_F/R[d] | GTTACGTGTGCAAAACTCATACCAGAATG | GATACTTGTTTAGGCTGAGAGTGCAGTGG |
| | | ACT2_F/R | GCACCCTGTTCTTCTTACCG | AACCCTCGTAGATTGGCACA |

[a-d]The sequences of these primers were taken from [a]Sarmiento-Mañús et al. (2023), [b]http://signal.salk.edu/tdnaprimers.2.html, [c]Garton et al. (2007), and [d]Rosa et al. (2013).
